# Predator prey and the third beneficiary

**DOI:** 10.1101/730895

**Authors:** Anagha Pund, Ketki Holkar, Milind Watve, Ulfat Baig

## Abstract

All bacterial epibiotic predators are rich in secondary metabolites and most genera rich in secondary metabolites have demonstrable predatory abilities. Therefore it is likely that an antibiotic resistant and thereby predation resistant species may benefit not only by escaping predation but also by utilizing nutrients released by lysis of prey cells by predatory bacteria. The resistant organisms may enjoy greater fitness benefits than the predator since they get the benefit without investing in the predation machinery. In our experiment, a marine isolate of *Streptomyces atrovirens* showed good predatory activity on a range of species including *Staphylococcus aureus* and *Proteus vulgaris. Escherichia coli* was resistant to predation by this species. On slide culture with water agar when the predator, *S. aureus* and *E. coli* were grown together *S. aureus* population declined whereas the predation resistant *E. coli* increased their population as compared to controls. However the growth of *E. coli* did not affect growth of the predator unfavorably. This strengthens the possibility that evolution of antibiotic resistance not only gave a selective advantage of escaping predation, it also would have increased the fitness of the resistant organism by promoting growth on nutrients released from the prey cells lysed by the predator. When the predator was grown with *S. aureus* and *P. vulgaris* as prey, *S. aureus* declined rapidly whereas *P. vulgaris* was spared. This suggests that the predator appears to show preference towards prey and in that case even a partial or relative resistance may give substantial advantage to a population.

## Introduction

Predation is defined as the act of consumption of a living organism by another living organism. Predation is a major ecological force, shaping the structure of communities, driving diversity and evolution of life histories (Stanley 1973; Day, Abrams and Chase 2002). Since bacteria do not have phagocytic abilities predatory bacteria lyse and degrade the prey cells using extracellular weapons including enzymes and secondary metabolites (Senges *et al.* 2018). Involvement of antibiotics and other secondary metabolites in predation is inferred from the strong association between predatory activity and genomic richness of secondary metabolites (Kumbhar and Watve 2013); differential expression of secondary metabolite genes when co-cultured with different prey species (Kumbhar *et al.* 2014) and loss of predatory abilities after mutating the antibiotic genes (Xiao *et al.* 2011). Kumbhar and Watve (2013) argued that antibiotics primarily evolved for predation, later diverging into mutualistic, signalling (Davies and Davies 2010) and other functions. Based on the association between antibiotics and predation Leisner et al (2016) argued that antibiotic resistance primarily evolved to resist predation but further this ability allows the resistant organisms to cross feed on the nutrients released from the lysis of susceptible prey cells by the predator. Resistant organisms can thus hitch hike on the predator and they might enjoy a fitness advantage over the predator since the predator needs to invest in the predation machinery which the hitch hiker doesn’t have to. Although this is a logical possibility, this has not been experimentally demonstrated so far.

*Streptomyces sp*. is known for producing an array of secondary metabolites (Watve *et al.* 2001). *Streptomyces spp.* are non-obligatory predators which are suspected to utilize a diversity of small molecules for predation (Kumbhar and Watve 2013; Kumbhar *et al.* 2014). The lysis of prey cells adjoining *Streptomyces* mycelium has been demonstrated microscopically using Differential Interference Contrast (DIC) microscopy (Kumbhar *et al.* 2014). In the present study we monitored the populations of susceptible and resistant prey species in presence of a predatory *Streptomyces*. We screened *S. atrovirens*, a predatory marine isolate against 14 lab strains and environmental isolates out of which 6 were susceptible to predation (*S.aureus, P.vulgaris, Mycobacterium smegmatis, Serratia marcencesus, Micrococcus leuteus and Bacillus subtilis*). After studying predatory activity against individual cultures we studied the responses of mixed populations of prey cells. The population response of a predation susceptible and resistant species, as well as two susceptible species co-cultured in presence of predator is reported here.

## Material and method

*Escherichia coli* (NCIM 2184), *Staphylococcus aureus* (NCIM 2121) and *Proteus vulgaris* (NCIM 2172) were inoculated in nutrient broth and incubated for 24 h at 37°C. Prey cells were centrifuged (Eppendorf centrifuge 5810R) at 6000 rpm for 10 min to concentrate cells followed by washing with sterile distilled water to remove traces of nutrients. After washing cells were suspended in sterile distilled water making thick slurry of cells. Prey cell number was standardized based upon a relationship between optical density of the washed suspension and the cell density obtained using Neubauer chamber (Rohem India BS 748). To study three species interaction we used two different co-culture combinations namely *E. coli* and *S. aureus* in approximately 1:1 ratio and *P. vulgaris* and *S. aureus* as prey in the same ratio. In these combinations it was possible to identify the cells based on morphology alone. It was not possible to study the *E. coli - P. vulgaris* combination owing to the difficulty of differential identification microscopically. The prey population was spread over the surface of water agarose bed on a slide culture over which *S. atrovirens* the predator, was spot inoculated. The agarose bed was covered with sterile coverslips so that same slide could be observed every day. The slides were incubated at 30°C for up to five days in moist chambers. Three controls were prepared similarly with (i) individual prey species in pure culture (ii) individual prey species in presence of predator and (iii) co-culture of two prey species without the predator.

Observations were made daily using DIC microscopy on Zeiss Axioimager M-1 upright microscope (100X/1.40 Oil DIC M27), alpha plan apochromat objective, with a digital camera, HAL 100 illuminator and quartz collector controlled by Axio software 4.8. DIC images were taken and resulting images were used for the quantification of cells. Ten randomly selected fields were imaged from every slide every day. The number of cells of each of the prey species per field was counted from the images. *S. atrovirens* being a mycelial species, we recorded in every image the number of times a diagonal crossed mycelial threads. The change in the population densities thus measured during the five day incubation period was studied.

## Reproducibility runs

Experiment was repeated three times at different time points (data not shown here) and similar trends were observed in all the repetitions. Since the starting populations were substantially different in every run we did not pool the results. In all the three experiments the results were qualitatively the same in that in every run, *S. aureus* population declined while *E. coli* population and the predator population increased in three species experiment. In case of three species experiment with two sensitive organisms *S.aureus* and *P. vulgaris* populations of both the organisms declined; *S. aureus* being first to show decline followed by decrease in *P. vulgaris* population. Predator, *S. atrovirens* increased in both the experiments.

## Results

Since the medium contained no nutrients, in pure culture controls the prey cells did not show a significant trend in population during the incubation period. In prey co-culture there was no significant trend in the populations of *S. aureus and E. coli* however *S. aureus and P. vulgaris* populations showed a growing trend when co-culturing indicating some synergistic interaction. We did not investigate the nature of this interaction. In one to one predator prey interactions *S. aureus and P. vulgaris* populations showed a monotonic declining trend while the predator population increased. In the *E. coli* predator interaction, neither the population of the predator nor that of *E. coli* showed a time trend.

In the three species interaction experiment with *S. aureus* and *E. coli* along with the predator, the *S. aureus* population declined by 97 % whereas the E. coli population increased by 60 % (Fig. 1C). The predator population increased by 72 % (Fig. 1D).

**Figure 1:**
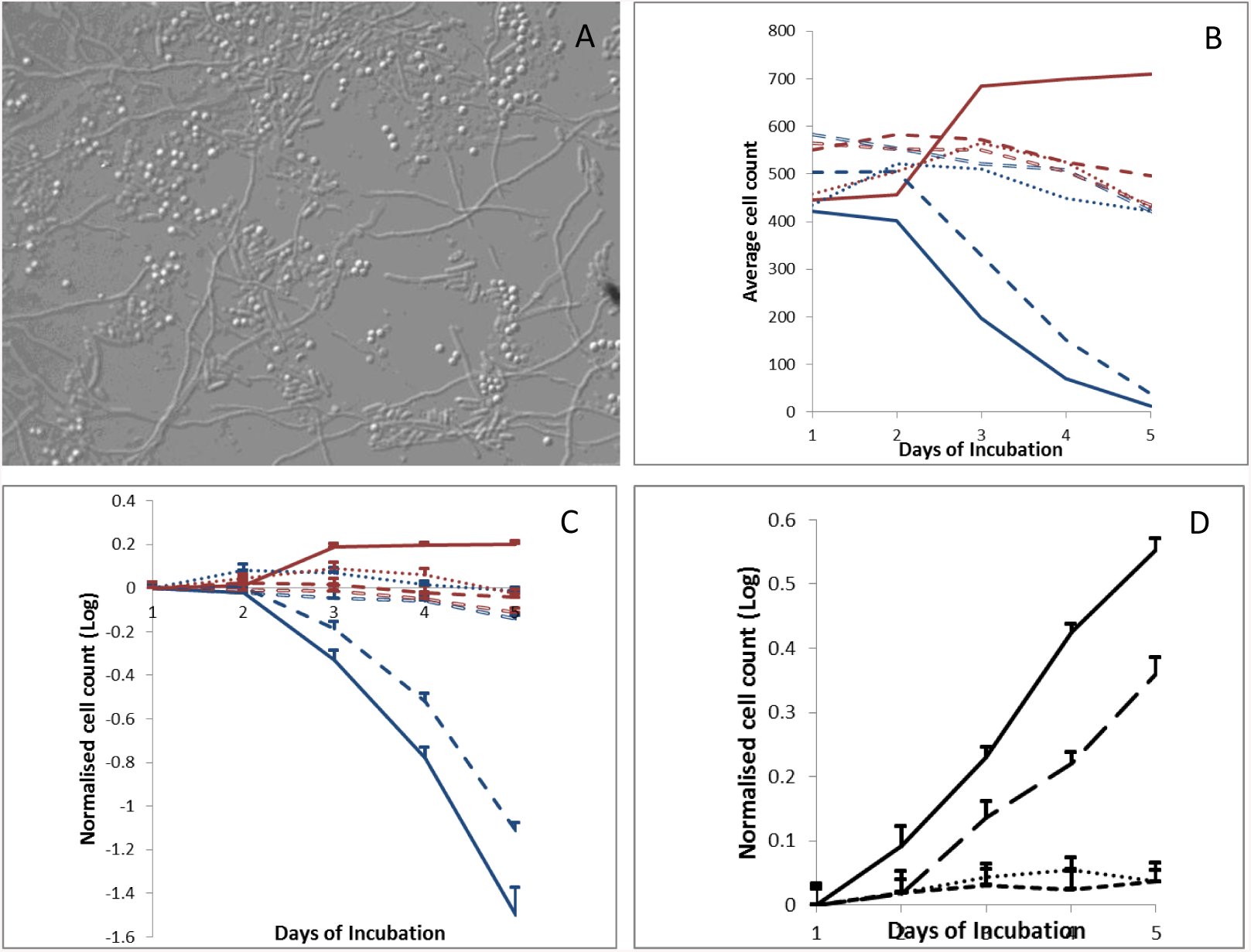
Change in cell density of prey predator and resistant cells in three species interaction. **A.** Microscopic image of slide culture for three species experiment with prey and resistant cells. **B.** Change in absolute average cell count of *S.aureus* (Solid blue line) and *E.coli* (Solid brown line) along with controls (*S.aureus* predation in solid dashed blue line, *S.aureus* in coculture with *E.coli* in hollow dashed blue line, pure *S.aureus* in dotted blue line and *E.coli* predation in solid dashed brown line, *E.coli* in coculture with *S.aureus* in hollow dashed brown line, pure *E.coli* in dotted brown line) over incubation period. **C.** Change in cell count of *S.aureus* and *E.coli* (normalized and on log scale) **D.** Change n cell count of predator, *S.atrovirens* (Solid black line) along with controls (*S.atrovirens* count in *S.aureus* predation, dashed black line, *S.atrovirens* count in *E.coli* predation, small dashed black line, pure *S.atrovirens* dotted black line.)

In the three species interaction experiment with *S. aureus* and *P. vulgaris* along with the predator by the second day, the *S. aureus* population declined by about 68% and remained low on the third day but *P. vulgaris* population did not show a significant change. However, during the last phase of growth the *P. vulgaris* population showed a decline by about 61% whereas *S. aureus* showed a small but significant increase (Fig. 2C). *S. atrovirens* population increased monotonically (Fig. 2D). This suggests that the predator might have selectively preferred *S. aureus* in the first phase and when its population declined below a threshold shifted to *P. vulgaris* allowing some comeback growth of *S. aureus*.

**Figure 2:**
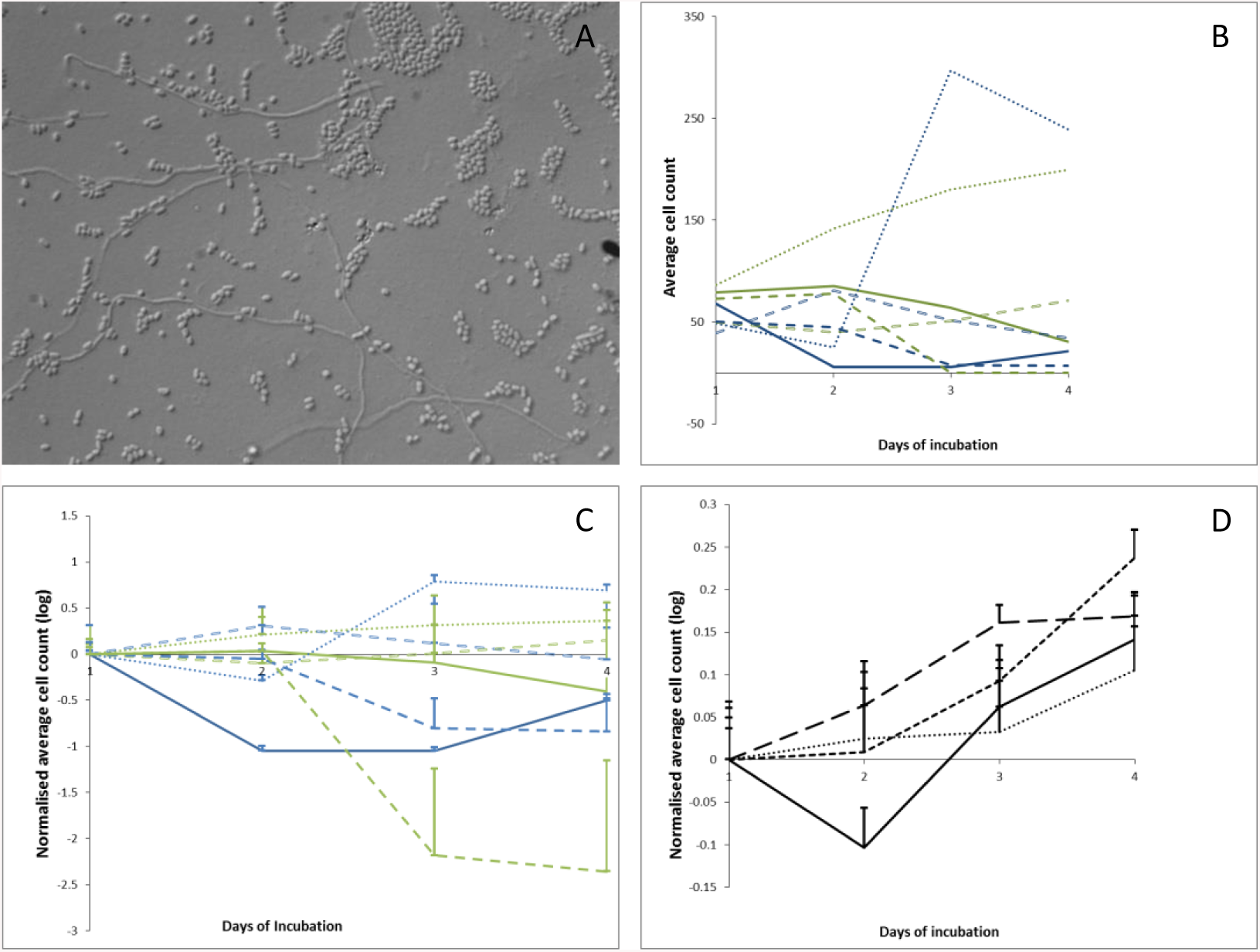
Change in cell density of predator and two sensitive prey cells in three species interaction. **A.** Microscopic image of slide culture for three species experiment with two sensitive cells. **B.** Change in absolute average cell count of *S.aureus* (Solid blue line) and *P. vulgaris* (Solid green line) along with controls (*S.aureus* predation in solid dashed blue line, *S.aureus* in coculture with *P. vulgaris* in dotted blue line, pure *S.aureus* in hollow dashed blue line and *P. vulgaris* predation in solid dashed green line, *P. vulgaris* in coculture with *S.aureus* in dotted green line, pure *P. vulgaris* in hollow dashed green line) over incubation period. **C.** Change in cell count of *S.aureus* and *P. vulgaris* (normalized and on log scale) **D.** Change in cell count of predator, *S.atrovirens* (Solid black line) along with controls (*S.atrovirens* count in *S.aureus* predation, long dashed black line, *S.atrovirens* count in *P. vulgaris* predation, small dashed black line, pure *S.atrovirens* dotted black line.)

## Discussion

The results highlight the complexity of multi-species interactions in bacteria. Earlier studies have shown that in the absence of other soluble nutrients, some bacterial genera turn predatory and grow at the expense of surrounding cells. We show here that different bacteria show differential sensitivity to predation which is important in shaping the patterns of interaction. Since bacterial predation is an extracellular phenomenon, a resistant organism gets an added nutritional benefit when in proximity of a predator and a susceptible organism. This was predicted earlier (Leisner, Jørgensen and Middelboe 2016) but our experiments have demonstrated this advantage of resistance empirically for the first time. The mechanism of predation resistance is not yet known. Secondary metabolites of the predator are suspected to have a role in predation (Kumbhar *et al.* 2014), therefore it is likely that resistance to the secondary metabolites might be the primary mechanism of predation resistance. If this is true it gives an added dimension to the evolution of antibiotic resistance.

More complex and more interesting appear to be the interaction between of two predation susceptible species. The predator that showed successful predation on both when tested separately appeared to exhibit preferential consumption of one species when challenged with both together. This appears to have spared the other species at least for some time. This demonstrates that even partial or relative resistance may give substantial selective advantages in natural multispecies settings.

Microbiology has largely progressed by pure culture studies. Natural ecosystems, on the other hand have a variety of multispecies interactions about which our understanding is still quite primitive. The experiments suggest that studying multispecies interactions will reveal a variety of novel dimensions of bacterial life in nature.

